# Hybrid zone analysis using coalescent-based estimates of introgression and migration

**DOI:** 10.1101/2025.01.21.634130

**Authors:** Adam D. Leaché, Hayden R. Davis, Sonal Singhal

## Abstract

Hybrid zones can serve as natural experiments allowing us to investigate the evolutionary consequences of introgression between distinct populations with divergent genomic backgrounds. Modeling hybrid zones using a coalescent framework can provide critical insights into the historical demography of populations, including population divergence times, population sizes, introgression proportions, migration rates, and the timing of hybrid zone formation. We used coalescent analyses to test whether a hybrid zone between Plateau Fence Lizard (*Sceloporus tristichus*) populations in Arizona formed recently because of human-induced landscape alterations or originated during Pleistocene climatic shifts. Introgression analysis places the divergence time between the parental populations at approximately 140 kya and their secondary contact and hybridization at approximately 10 kya at the end of the Pleistocene. Introgression proportions for hybrid populations are correlated with their geographic distance from the parental populations, a pattern that is repeated with admixture proportions and hybrid index values. The multispecies coalescent with migration model supports asymmetric migration rates with a bias for gene flow moving towards the northern end of the hybrid zone. The direction of this asymmetry is consistent with spatial cline analyses that suggest a slow but steady northward shift of the center of the hybrid zone. When analyzing hybrid populations sampled along a linear transect, coalescent methods can provide novel insights into the evolutionary history of hybrid zone formation that complement existing spatial and genomic methods.

## Introduction

Hybrid zones are geographic areas where genetically distinct populations produce offspring with a mixture of traits from the parental populations (Harrison 1993; Gompert *et al*. 2017). Naturally occurring hybrid zones provide important insights into the evolutionary processes of speciation, gene flow, and species boundaries (Harrison & Larson 2014). Spatial clines and genomic clines are two standard approaches used to study hybrid zones. Spatial clines are important for understanding geographic characteristics of a hybrid zone, including the location of the center of the hybrid zone (cline center), the width of the hybrid zone (cline width), and when multiple loci are compared, differential introgression among marker types (Barton & Hewitt 1985; Szymura & Barton 1986). Genomic clines leverage large numbers of loci from across the genome to provide insights into differential introgression among genes or chromosomal regions (Gompert & Buerkle 2011). Hybrid zones can also be modeled using a coalescent framework to investigate the historical demography of populations, which adds critical information on divergence times, population sizes, introgression rates, the timing of hybrid zone formation, and the intensity of introgression from parental to hybrid populations (Jiao *et al*. 2021; Ji *et al*. 2023). In this study, we use coalescent models to test hypotheses regarding the age of a hybrid zone and extent of gene flow between populations of the Plateau Fence Lizard (*Sceloporus tristichus*) in southwestern North America.

The *Sceloporus tristichus* hybrid zone is in Arizona’s Colorado Plateau at an ecological transition zone between Great Basin Conifer Woodlands in the south and Grassland habitats in the north; this transition zone facilitates hybridization between morphologically and ecologically distinctive grassland and woodland populations (Leaché & Cole 2007). The canyon ecotone habitats that support hybridization may have formed as recently as the 1890s with the onset of intensive cattle grazing, which mediated the northward expansion of juniper trees into former grasslands (Archer 1994; Abruzzi 1995). Alternatively, palynological evidence suggests that juniper expansion in the American Southwest began at least several thousand years earlier (Davis *&* Turner 1986; Miller *&* Wigand 1994). To test these alternative hypotheses explaining the formation of the hybrid zone we use the multispecies coalescent with introgression (MSC-I) model, which provides a natural way to estimate the introgression time between populations (Fig. 1).

**Figure 1.**
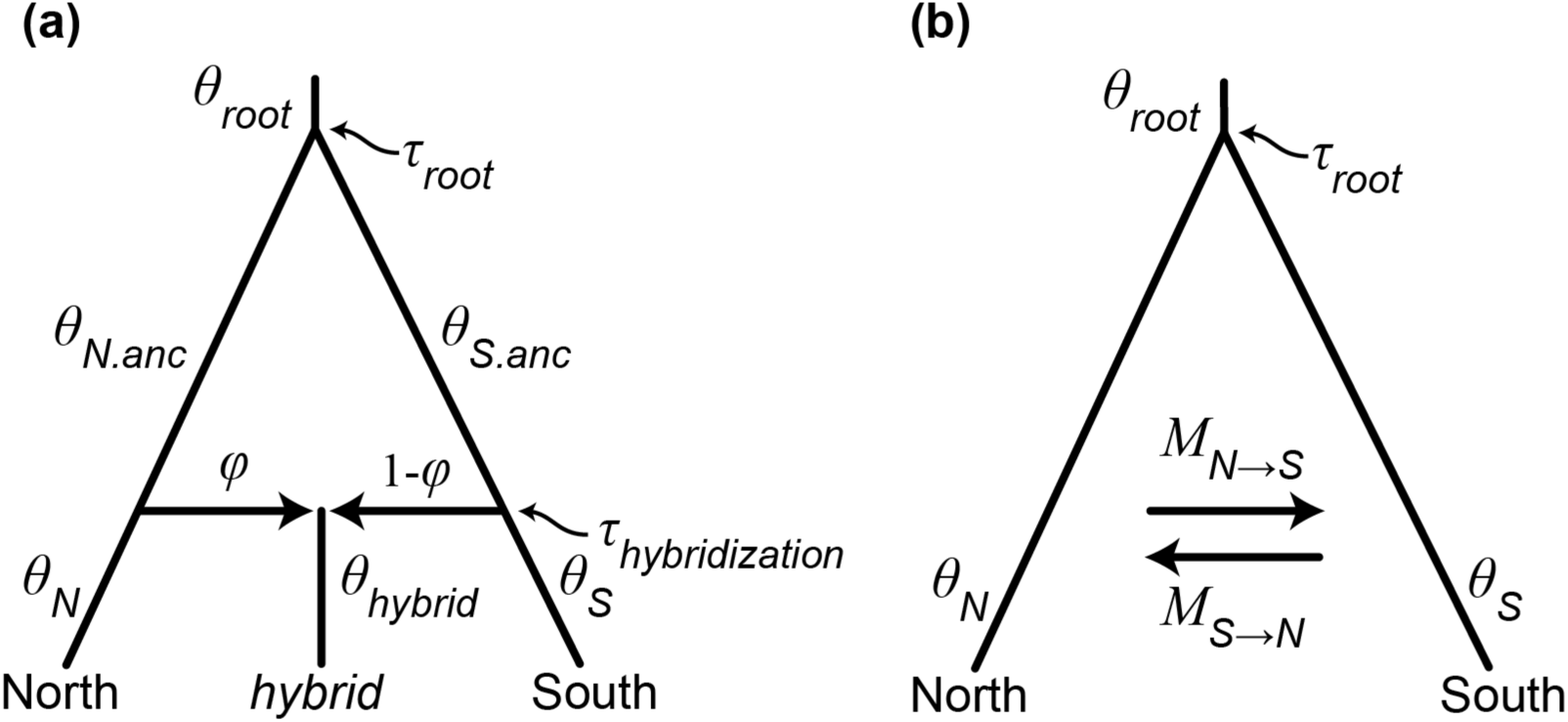
Multispecies coalescent models used to estimate population demographic parameters in the hybrid zone. (a) The introgression model (MSC-I) includes nine parameters: six population sizes (*θ_N_, θ_S_, θ_hybrid_, θ_N.anc_, θ_S.anc_, θ_root_*), one divergence time (*τ*_root_), one hybridization time (*τ*_hybridization_), and one introgression probability (*φ*) defined as the proportion of the hybrid population genome originating from one of the parental populations (the proportion of the genome originating from the other population is 1-*φ*). Parameters for the MSC-I model were estimated separately for each hybrid population. (b) The migration model (MSC-M) includes six parameters: three population sizes (*θ_N_, θ_S_, θ_root_*), one divergence time (*τ*_root_), and two migration rates (*M_N→S_*, *M_S→N_*) defined as the expected number of migrants from the donor population to the recipient population per generation.

Comparisons of spatial clines using hybrid zone samples collected over multiple decades have provided evidence that the center of the cline is moving to the north (Leaché *&* Cole 2007; Leaché *et al* 2017). Specimens collected in the 1970s and 2002 show changes in the frequency of a chromosome inversion polymorphism (e.g., a distinctive pericentric inversion polymorphism on chromosome seven) that is consistent with the hypothesis of northward hybrid zone movement (Leaché & Cole 2007). Also, DNA sequence comparisons of samples collected in 2002 and 2012 provide evidence that the mitochondrial DNA (mtDNA) cline is displaced from the nuclear cline and maintaining an introgression distance of approximately 3 km, and both clines shifted north by approximately 2 km over this time-period, which equals roughly 10 lizard generations (Leaché *et al*. 2017). In this study, we use a new replicate of transect sample from 2022 to test the prediction that the hybrid zone has continued to shift to the north using spatial cline analyses. Furthermore, we use the MSC with migration (MSC-M) model to estimate bi-directional migration rates between the parental populations, which could show a bias towards northward migration if the hybrid zone has moved through time.

## Materials and Methods

### Sampling

We collected 79 samples from eight localities spanning the hybrid zone in 2022 (Table 1; Table S1, Supporting information). The sampling transect extends 63.5 km from grassland habitat in the north to juniper woodland habitat in the south. Two localities (Holbrook and Show Low) are parental populations at the tails of the hybrid zone, and the other six populations constitute the transect through the ecotone. Canyon ecotones that connect these habitats are where lizards with mixed genotypes are found (Leaché & Cole 2007). We compared the 2022 transect to 95 samples collected in 2002 (Leaché & Cole 2007) and 179 samples collected in 2012 (Leaché *et al*. 2017) from the same localities. Scientific collecting was approved by the State of Arizona Game and Fish Department (SP# SP843452). Animal research was approved by the University of Washington Office of Animal Welfare (IACUC #4367-03).

**Table 1.** Hybrid zone transect sites (*n* = 8) compared over 10-year intervals using replicate sampling. Distance (km) is from the northernmost locality (Holbrook). Sample sizes (*N*) for genomic DNA sequencing are shown for each transect year. The ancestry proportions (*Q*) estimated using ADMIXTURE are shown as the fraction of ancestry to the northern population under a *K* = 2 model.

|  |  | 2002 |  | 2012 |  | 2022 |  | All years |  |
| --- | --- | --- | --- | --- | --- | --- | --- | --- | --- |
| Site name | Distance | $N$ | $Q$ | $N$ | $Q$ | $N$ | $Q$ | $N$ | $Q$ |
| Holbrook | 0 | 9 | 0.98 | 20 | 1.00 | 7 | 1.00 | 36 | 0.99 |
| Fivemile Wash | 8.3 | 8 | 1.00 | 20 | 0.89 | 11 | 0.99 | 39 | 0.92 |
| Washboard Wash | 14.5 | 9 | 1.00 | 20 | 0.98 | 10 | 0.99 | 39 | 0.98 |
| Woodruff | 22.6 | 4 | 1.00 | 20 | 0.98 | 9 | 1.00 | 33 | 0.98 |
| Canoncito | 29.3 | 24 | 0.89 | 40 | 0.83 | 10 | 0.87 | 74 | 0.81 |
| Sevenmile Draw | 35.2 | 8 | 0.29 | 20 | 0.23 | 10 | 0.22 | 38 | 0.25 |
| Snowflake | 42.7 | 7 | 0.00 | 19 | 0.02 | 10 | 0.01 | 36 | 0.05 |
| Show Low | 63.5 | 9 | 0.00 | 20 | 0.00 | 11 | 0.00 | 40 | 0.0 |

### Mitochondrial DNA analysis

Mitochondrial DNA (mtDNA) haplotypes from two different species, *Sceloporus tristichus* and *S. cowlesi* are found in the hybrid zone (Fig. 2), and the *S. tristichus* haplotypes belong to three divergent clades (Leaché & Cole 2007). Previous analyses of mtDNA in the hybrid zone have detected all four mtDNA haplotypes in the center of the hybrid zone (Leaché and Cole 2007). The four mtDNA haplotype clades found in the hybrid zone are as follows: (1) *S. tristichus* north is a clade containing northern hybrid zone populations; (2) *S. tristichus* south is a clade contains southern hybrid zone populations; (3) *S. tristichus* west is a clade that leaks into the hybrid zone from the west into several hybrid populations; and (4) *S. cowlesi* is a separate species with mtDNA haplotypes that enter the hybrid zone from the east and are found in the hybrid populations (Fig. 2).

**Figure 2.**
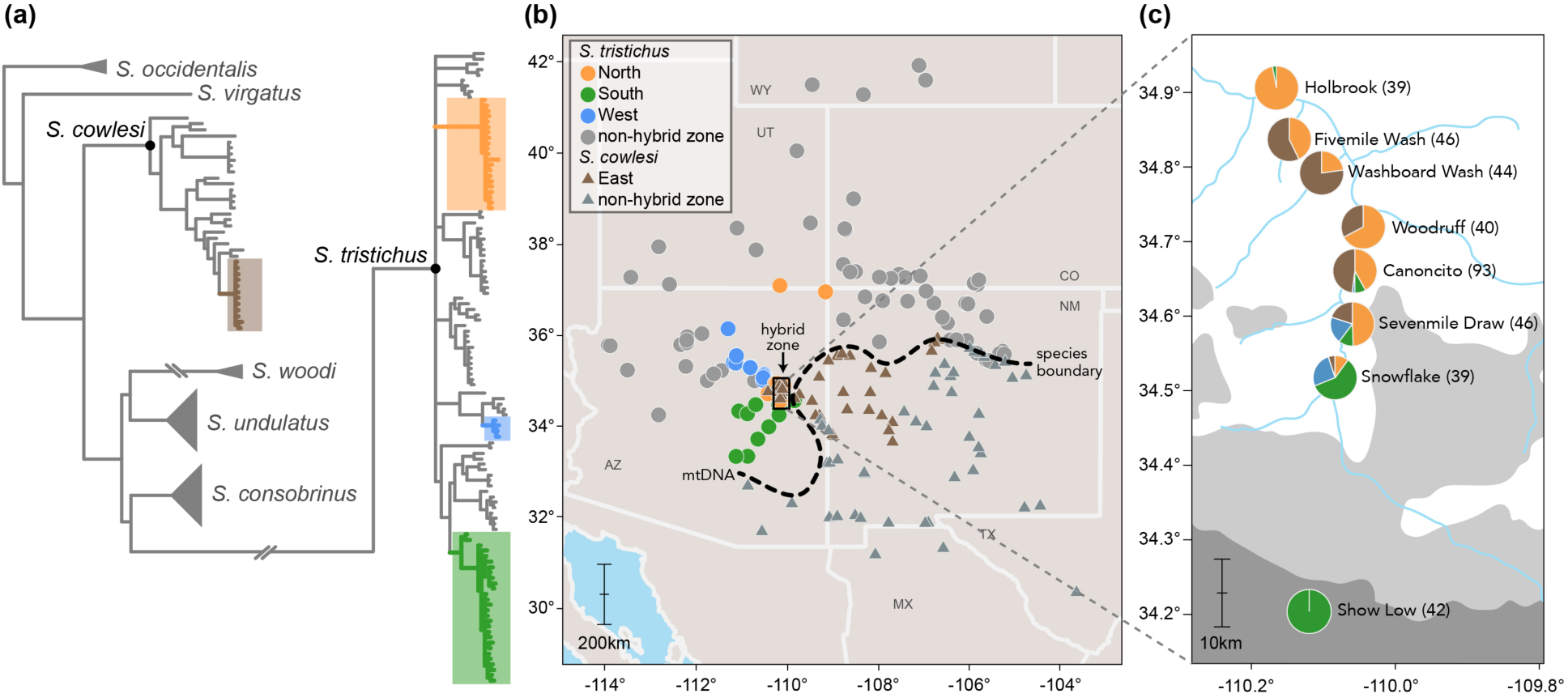
Phylogeny and geographic distributions of the study species, *Sceloporus tristichus* and *S. cowlesi* in the Southwestern US. (a) Phylogenetic relationships based on mitochondrial DNA (mtDNA) sequence data (*ND1* gene) estimated using maximum likelihood. The four mtDNA clades that are found in the hybrid zone are color-coded. (b) Geographic distributions of *S. tristichus* (circles) and *S. cowlesi* (triangles) based on mtDNA sequence data, the distributions of the clades detected in the hybrid zone, and the mtDNA species boundary (dashed line). (c) Pie-charts showing mtDNA haplotype frequencies in the hybrid zone. Sample sizes are shown in parentheses. The shaded areas show habitat distributions of Petran Montane Conifer Forest (dark grey), Great Basin Conifer Woodland (light grey), and Great Basin Grassland (unshaded/white).

For both the genomic and mtDNA sequencing, we extracted genomic DNA from tissue samples using salt extraction (Aljanabi & Martinez 1997). To determine the frequency of haplotypes at each hybrid zone sample site, we amplified and sequenced the mitochondrial *ND1* protein-coding gene (969 bp) for the 2022 samples using standard PCR methods (Leaché & Cole 2007). In addition, we expanded our sampling of *S. cowlesi* and *S. tristichus* to update and refine the geographic boundaries between these mtDNA-defined species. We aligned the new *ND1* sequences with previously published sequences (Leaché *et al*. 2017), which included other closely related *Sceloporus* species (e.g., *S. cautus, S. consobrinus, S. exsul, S. occidentalis, S. undulatus, S. virgatus,* and *S. woodi*). New mtDNA sequences are deposited in GenBank (Accession nos XXXXXXXX–XXXXXXXX), and the alignment is available on Dryad (doi:XX.XXXX/dryad.XXXX). We estimated a phylogeny using maximum likelihood using RAxML-NG v.1.1.0 (Kozlov *et al*. 2019) with the GTR+GAMMA substitution model (following previous studies of the hybrid zone) and 100 bootstrap replicates.

### SNP data collection

We collected SNP data using the double digestion restriction site associated DNA sequencing (ddRADseq) protocol (Peterson *et al*. 2012). We followed the same genomic library preparation methods used in a previous study (Leaché *et al*. 2017). Briefly, we double-digested genomic DNA with enzymes SbfI and MspI, ligated barcodes, adapters, unique molecular identifiers (UMIs), and then pooled sampled and size-selected, and PCR amplified the libraries. Samples were sequenced on one lane (100bp single-end reads) with an Illumina NovaSeq 6000 at the QB3 facility at UC Berkeley.

### Bioinformatics

We processed raw Illumina reads using the program ipyrad v.0.7.3 (Eaton & Overcast 2020). We de-multiplexed samples using their unique barcode and adapter sequences, and sites with Phred quality scores under 99% (Phred score = 20) were changed into “N” characters and reads with ≥ 10% N’s were discarded. After the removal of the 6 bp restriction site overhang, the 6 bp barcode, and 8 bp UMI, all reads were trimmed from the 3’ end to match the length of reads collected previously (39bp). We aligned reads using a reference genome of *Sceloporus tristichus* collected from the northern end of the hybrid zone (Holbrook; Bedoya & Leaché 2021) using a clustering threshold of 90%. Additional filtering included discarding clusters that had low coverage (< 10 reads), excessive undetermined or heterozygous sites (> 4), or too many haplotypes (> 2 for diploids). The final datasets included loci with no more than 50% missing data. Additional filtering was conducted using VCFtools v.0.1.16 (Danecek *et al*. 2011) to sample biallelic loci, impose a minor allele frequency threshold of 0.01, and to retain one SNP per locus.

To assess the robustness of the cline fitting analyses, we produced two different assemblies. The combined assembly included all 335 hybrid zone samples collected from 2002, 2012, and 2022. This assembly maximized the number of loci that overlap across transect time-series, but since it includes the most samples it also increases missing data. We also produced separate assemblies for the transect sampling years (2002, 2012, 2022). These assemblies maximize the number of loci obtained for each sampling year and minimize missing data compared to the combined assembly. De-multiplexed sequences are available on the NCBI Sequence Read Archive (PRJNA1211730), and the final assemblies are available on Dryad (doi:XX.XXXX/dryad.XXXX).

### Population structure

We estimated population structure using the SNP data with ADMIXTURE v.1.3.0 (Alexander *et al*. 2009). The hybrid zone contains two well-defined genetic clusters (Leaché *et al*. 2017), so we used *K* = 2 to estimate ancestry proportions for individuals in each of the 8 localities. Each analysis was run 10 times using random starting seeds. The admixture proportions under the *K* = 2 model represent the fraction of membership of each locality to the northern and southern parental populations.

### Cline-fitting analysis

We performed ML cline-fitting analyses of the SNP data assemblies using the R package hzar (Derryberry *et al*. 2014). Clines were fit assuming allele frequency intervals fixed at 0 and 1 without tail fitting (Szymura & Barton 1986; Szymura 1991). Our previous study of the hybrid zone found this model to be a better fit compared to three alternatives using AIC model selection (Leaché *et al*. 2017). We used the ancestry proportion estimates from ADMIXTURE for maximum likelihood (ML) cline model inference. We estimated ML clines for each sampling period (2002, 2012, 2022) using the combined and separate data assemblies. Cline parameters of interest included cline center (measured as distance in kilometers from the northern parental population) and cline width (kilometers). Tests for parameter uncertainty were estimated using 2-log-likelihood (2LL) support limits.

### Hybrid index analysis

We used the R package gghybrid (Bailey 2024) to estimate the genome-wide hybrid index *h* of each hybrid sample. The hybrid index represents the proportion of the genome originating from parental reference populations; Holbrook (northern) and Show Low (southern) (Table 1). Under this model, each allele is treated independently, and the probability that an individual allele originated from one of the source populations depends on the frequency of the allele in each source population and the hybrid index of the test subject (Buerkle 2005). We followed the recommended analysis procedures and ran the chain for 3000 iterations with a 1000 step burn-in period (Bailey 2024).

### Multispecies coalescent with introgression or migration

We used the multispecies coalescent with introgression (MSC-I) model in the program BPP v.4.7 (Rannala & Yang 2003; Flouri *et al*. 2020) to estimate the timing of hybridization and the intensity of introgression from each parental population into the hybrid zone. The hybrid zone model includes three populations; two parental populations that diverge at *τ*_root_ and later hybridize at time *τ*_hybridization_ leading to a hybrid population (Fig. 1a). This model is described as a hybrid speciation model whereby populations come into contact to form a hybrid species (Flouri *et al*. 2020), and we consider this model a good representation for the *Sceloporus* hybrid zone where two populations have come into contact to form a hybrid population. The introgression probability (*φ*) is the proportion of the hybrid population genome that is derived from the parent population. A limitation of this model is that it does not account for any introgression that occurs between the initial hybridization event and the present day, nor for potential introgression from unsampled populations (here, other hybrid zone localities).

We conducted six separate MSC-I analyses to estimate *τ*_hybridization_ and *φ*_North→*x*_ for each of the hybrid populations while holding the parental populations constant (north = Holbrook; south = Show Low). These analyses used the new 2022 transect data assembled with no missing data (*n*=738 loci), since this dataset contained the most loci. Allowing no missing data ensured that the results using different hybrid populations were comparable by maintaining the same loci across all six analyses. We used gamma priors for population sizes (*θ*) and the root age (*τ*_root_), with the prior mean G(2, 1000) = 0.002, which are close to empirical estimates derived from the study system. The prior for introgression probability (*φ*) used a beta distribution ∼ beta(1,1), which provides a flat prior distribution. Analyses were run for 400,000 steps (sampling every other step) after a burn-in period of 40,000 steps. Convergence was assessed by running the same analysis four times with different starting seeds and confirming that the posterior distributions for parameters were similar between separate analyses (Flouri *et al*. 2018). The hybridization time (*τ*_hybridization_) and root age (*τ*_root_) were converted to absolute time assuming a lizard-specific substitution rate of 3.17^-9^ mutations/site/year (Bergeron *et al*. 2023) and a one-year generation time (Tinkle & Dunham 1986).

To estimate the level of gene flow between the parental populations without the influence of the hybrid samples, we estimated bi-directional migration rates using the MSC with migration model (MSC-M; Flouri *et al*. 2023). The simple two-population model used here includes six parameters: two migration rates, three population sizes (two populations and their ancestor), and one divergence time (Fig. 1b). The migration rate *M*=*mN* is measured in the expected number of migrants from the donor population to the recipient population per generation; *N* is the (effective) population size of the recipient population, and *m* is the proportion of immigrants in the recipient population (from the donor population) every generation, with time moving forward (Flouri et al., 2023). The migration rate prior used a gamma distribution G(1,10) with mean 1/10=0.1 migrant individuals per generation (1 migrant individual every 10 generations). All other priors and settings matched those used in the MSC-I analyses described above.

## Results

### Mitochondrial DNA analysis

We conducted a phylogenetic analysis of the *ND1* gene to calculate the frequency of mtDNA haplotypes found at each site in the hybrid zone. The mtDNA phylogeny places the hybrid zone samples into the four distinct clades defined in previous studies; three of these clades belong to *S. tristichus* and one belongs to *S. cowelsi* (Fig. 2). The northern *S. tristichus* haplotypes are found in high frequency (97%) in the northernmost population (Holbrook) and decline to 0% in the south (Show Low), while the southern *S. tristichus* haplotypes show the opposite pattern (Table 2). The western *S. tristichus* haplotypes do not form a cline, but instead are found at relatively low frequency (2% to 26%) in three of the southern populations. The *S. cowlesi* haplotypes enter the hybrid zone from the east and are found in the middle of the hybrid zone from Fivemile Wash to Snowflake and are in high frequency in the northern populations (57% and 77%; Table 2). Haplotypes belonging to all four mtDNA clades are found in three populations: Canoncito, Sevenmile Draw, and Snowflake (Table 2).

**Table 2.**
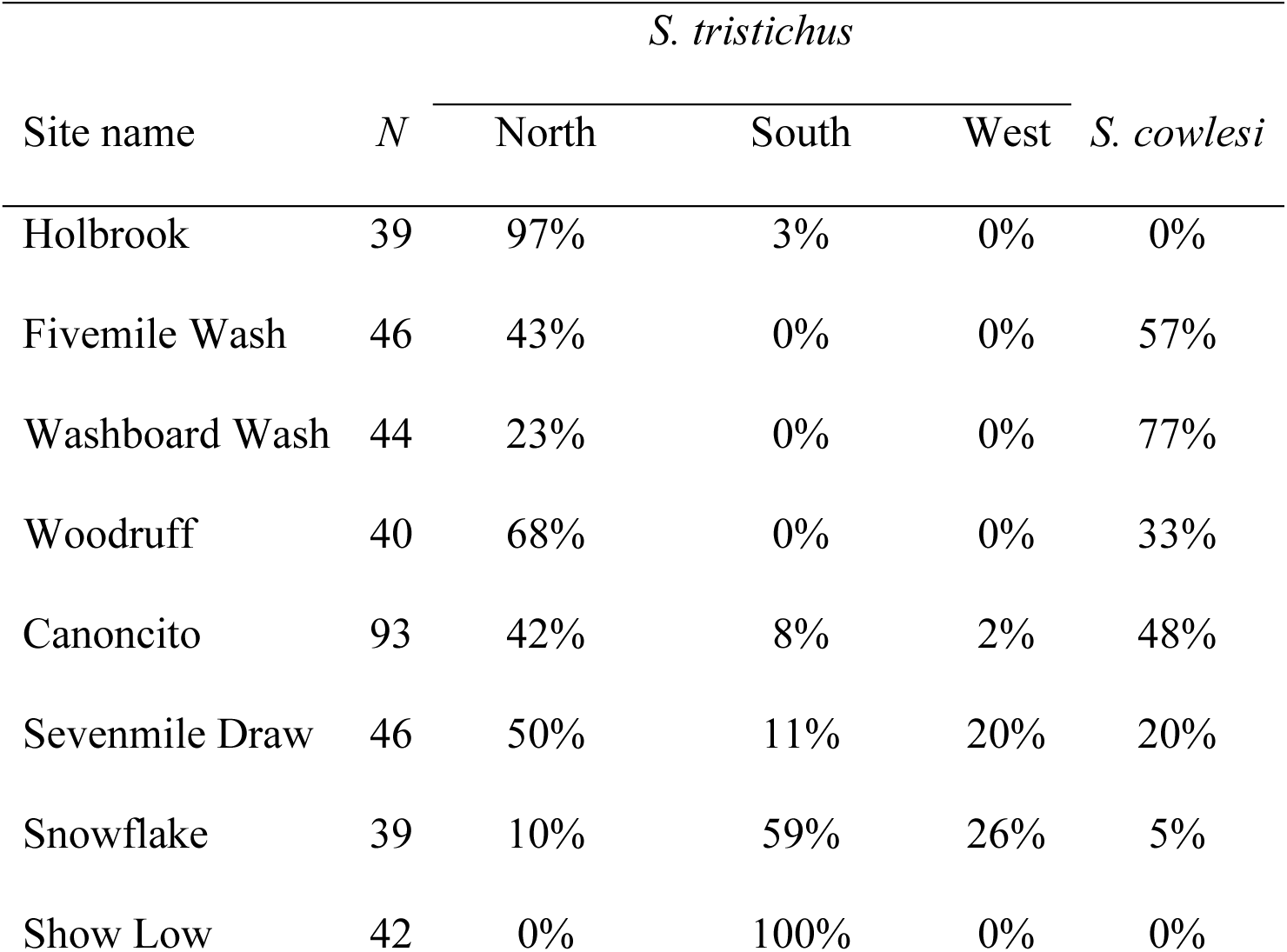
Frequency of *Sceloporus tristichus* and *S. cowlesi* mtDNA haplotypes at hybrid zone localities aggregated across three distinct time points (2002, 2012, and 2022). Haplotypes from *S. cowlesi* enter the hybrid zone from the east and are found at all sites except for the parental populations in Holbrook and Show Low.

### SNP data

The ipyrad reference assembly for the 2022 hybrid zone transect samples produced 3,803 ddRADseq loci after filtering for biallelic sites, a minor allele frequency of 0.01, and a missing data threshold of 0.5 (Table 3, Table S2, Supporting information). The separate reference assemblies for the 2002 and 2012 transect produced 1,872 and 1,171 loci, respectively (Table 3). Combining all samples together into one assembly produced fewer loci overall (2002 = 1,713 loci; 2012 = 1,137 loci; 2022 = 1,808 loci).

**Table 3.**
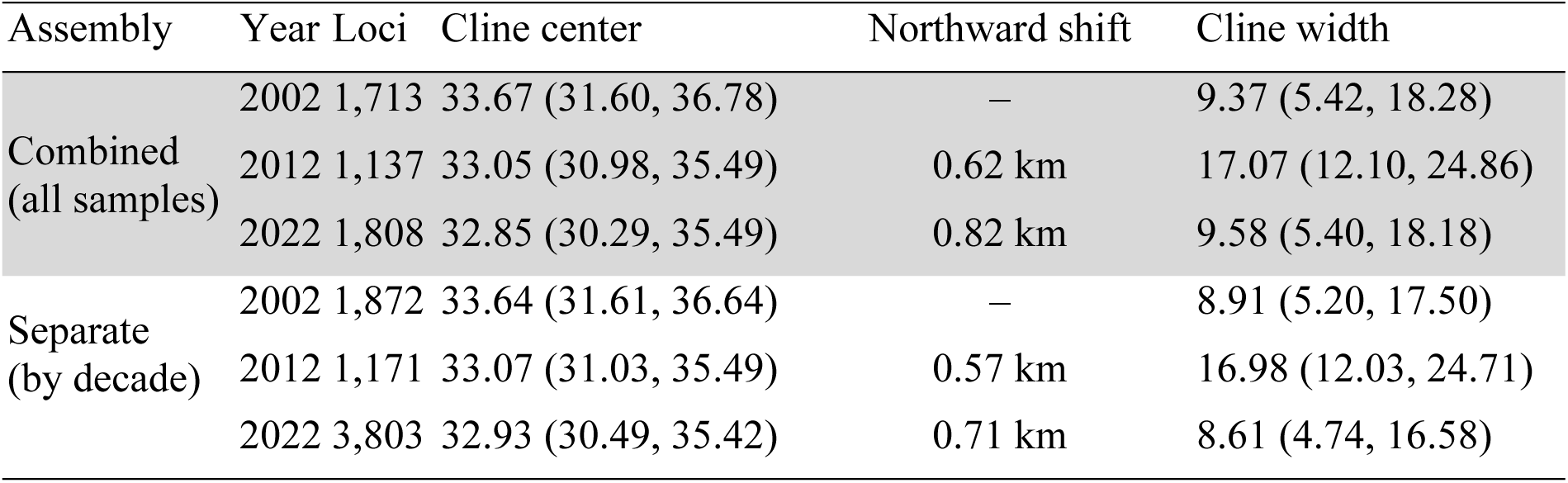
Temporal comparison of maximum likelihood cline centers and widths for the *Sceloporus tristichus* hybrid zone. The northward shift (compared to 2002) in the cline center is consistent across years and data assemblies. Distances are in units of kilometers, and the 2-log-likelihood support limits (2LL) are in parentheses (minimum, maximum).

| Assembly | Year | Loci | Cline center | Northward shift | Cline width |
| --- | --- | --- | --- | --- | --- |
| Combined<br>(all samples) | 2002 | 1,713 | 33.67 (31.60, 36.78) | – | 9.37 (5.42, 18.28) |
|  | 2012 | 1,137 | 33.05 (30.98, 35.49) | 0.62 km | 17.07 (12.10, 24.86) |
|  | 2022 | 1,808 | 32.85 (30.29, 35.49) | 0.82 km | 9.58 (5.40, 18.18) |
| Separate<br>(by decade) | 2002 | 1,872 | 33.64 (31.61, 36.64) | – | 8.91 (5.20, 17.50) |
|  | 2012 | 1,171 | 33.07 (31.03, 35.49) | 0.57 km | 16.98 (12.03, 24.71) |
|  | 2022 | 3,803 | 32.93 (30.49, 35.42) | 0.71 km | 8.61 (4.74, 16.58) |

### Population structure

Population structure estimates under a *K* = 2 model using the SNP data are shown for each hybrid zone location in Table 1. For each temporal transect, the ancestry proportions show a steep transition from northern ancestry to southern ancestry in the center of the hybrid zone (Table 1).

### Cline-fitting analysis

Spatial cline-fitting analyses of the 2022 SNP data were conducted to test the prediction that the hybrid zone is shifting to the north by comparing ML cline center estimates between decades. Movement of the hybrid zone is supported by the northward shift in cline centers, and this result is robust to the different data assemblies tested (Table 3). However, although the ML cline center estimates show directional movement, the magnitude of change is not significant when considering the overlap in 2LL support limits (Table 3). Cline width estimates range from approximately 9–17 km (Table 3).

### Hybrid index analysis

To calculate the proportion of the genome originating from each of the parental populations, we estimated the hybrid index *h* of each sample. The hybrid index plot supports a transition from northern to southern ancestry with greater northern ancestry in Woodruff, Washboard Wash, Fivemile Wash, and Canoncito, and greater southern ancestry in Sevenmile Draw and Snowflake (Fig. 3a). A histogram of hybrid index values shows a deficit of F1 hybrids at *h* = 0.5, and instead suggests that most samples represent late-stage backcrosses (Fig. 3b).

**Figure 3.**
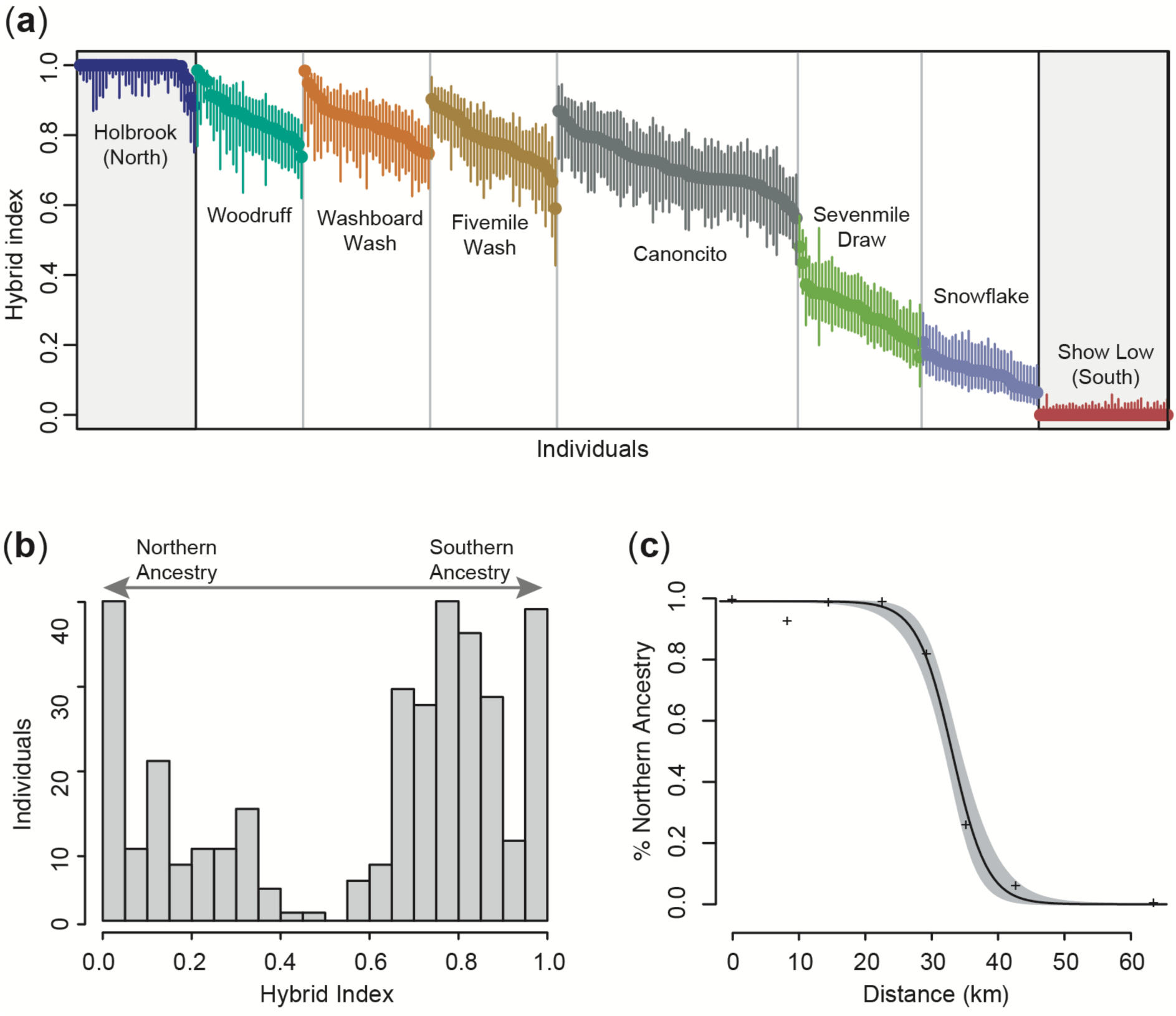
(a) Hybrid index plot for the *Sceloporus tristichus* hybrid zone. The hybrid index (estimated using gghybrid) is plotted for each individual (posterior mode) and error bars are 95% credible intervals. The points are ordered by population mean and then individual estimates. (b) Histogram of hybrid index values shows a high frequency of late-stage backcrosses with northern and southern parental populations. (c) Maximum likelihood cline for all samples combined. The ML cline is in black, confidence interval is shaded in grey, and observed admixture proportions are shown with “+” symbols.

### Multispecies coalescent with introgression or migration

To estimate the timing of hybridization and the intensity of introgression from the parental populations into the hybrid zone, we conducted six separate analyses using the MSC-I model (Fig. 1a). Estimates of the introgression probability (*φ*) from the northern population (Holbrook) into each hybrid population decline as the geographic distance from the northern population increases; conversely, *φ* from the southern population (Show Low) into each hybrid population increases along this same gradient. The estimates for *φ*_North→*x*_ range from ∼0.81 in the northern hybrid zone locations to as low as ∼0.12 in the south (Table 4). The MSC-I model contains two parameters related to time (Fig. 1a), including the divergence time of the parental populations (*τ*_root_) and hybridization event (*τ*_hybridization_). Estimates for *τ*_root_ vary for each analysis, but generally overlap with a range from 138–153 kya with the 95% highest posterior density (HPD) including 114–181 kya (Table 4). The estimates for *τ*_hybridization_ are an order of magnitude younger with the most recent estimate at 6.98 kya (2.8– 12.6 kya 95% HPD) for the population closest to the center of the hybrid zone (Canoncito; Table 4). A southern population, Sevenmile Draw, provides the oldest *τ*_hybridization_ = 16.03 kya (4.1–25.2 kya 95% HPD).

**Table 4.** Parameter estimates from the multispecies coalescent with introgression (MSC-I) model for the *Sceloporus tristichus* hybrid zone. For each hybrid population (*x*), the introgression probability (*φ*_North→*x*_) is the proportion of the hybrid population genome originating from the northern parental population (Holbrook). The proportion of the genome originating from the southern parental population (*φ*_South→*x*_) is 1-*φ*_North→ *x*_ (not shown). The hybridization time (*τ*_hybridization_) and root age (*τ*_root_) are converted to units of thousands of years (kya) assuming a lizard-specific substitution rate of 3.17^-9^ mutations/site/year. Bayesian posterior distributions are summarized by their mean and 95% HPD (in parentheses).

| Hybrid pop ($x$) | $\phi_{\text{North} \rightarrow x}$ | $\tau_{\text{hybridization}}$ | $\tau_{\text{root}}$ |
| --- | --- | --- | --- |
| Fivemile Wash | 0.729 (0.65–0.81) | 10.32 (3.5–14.2) | 144.40 (116.1–171.0) |
| Washboard Wash | 0.807 (0.72–0.88) | 11.96 (6.3–12.9) | 145.19 (118.9–171.6) |
| Woodruff | 0.748 (0.67–0.83) | 11.95 (6.6–17.4) | 143.11 (114.2–168.8) |
| Canoncito | 0.649 (0.57–0.73) | 6.98 (2.8–12.6) | 138.47 (116.7–161.2) |
| Sevenmile Draw | 0.299 (0.21–0.39) | 16.03 (4.1–25.2) | 151.78 (122.7–181.1) |
| Snowflake | 0.121 (0.06–0.18) | 11.27 (4.7–17.7) | 153.03 (128.7–178.2) |

To estimate the migration rate between the parental populations, we used the MSC-M model (Fig. 1b). The migration rates (*M*) are asymmetric with a bias from south to north with approximately 0.3 migrants per generation, or 3 migrants every 10 generations (*M* _South→North_ = 0.315; 95% HPD = 0.179–0.445; Table 5). Migration in the opposite direction is an order of magnitude lower and with a 95% highest posterior density (95% HPD) that overlaps with 0.0, suggesting that migration from north to south is not significant (*M* _North→South_ = 0.033; 95% HPD = 0.000–0.099; Table 5). The divergence time estimated under the MSC-M model is *τ*_root_ = 244.5 kya (95% HPD = 163.4–334.38 kya; Table 5), which is older but overlapping with the estimates from the MSC-I models (Table 4).

**Table 5.** Parameter estimates from the multispecies coalescent with migration (MSC-M) model for the parental populations (Holbrook, north and Show Low, south) in the *Sceloporus tristichus* hybrid zone. The migration rate (*M*) is the expected number of migrants from the donor population to the recipient population per generation. The divergence time (*τ*_root_) is converted to units of thousands of years (kya) assuming a lizard-specific substitution rate of 3.17^-9^ mutations/site/year.

| Parameter | Mean | 95% HPD |
| --- | --- | --- |
| $\theta_{\text{North}}$ | 0.00181 | 0.00132–0.00234 |
| $\theta_{\text{South}}$ | 0.00434 | 0.00344–0.00529 |
| $\theta_{\text{Ancestor}}$ | 0.00166 | 0.00113–0.00219 |
| $\tau_{\text{root}}$ | 244.5 kya | 163.4–334.38 kya |
| $M_{\text{North} \rightarrow \text{South}}$ | 0.03313 | 0.000–0.099 |
| $M_{\text{South} \rightarrow \text{North}}$ | 0.315 | 0.179–0.445 |

## Discussion

### Insights from spatial, genomic, coalescent approaches

Analyses of the *Sceloporus tristichus* hybrid zone using spatial, genomic, and coalescent approaches provide complementary inferences regarding the clinal nature of the hybrid zone. Population structure analyses of the hybrid zone show a sharp transition in ancestry proportions in the middle of the hybrid zone (Table 1). Analyzing these data in a spatial framework with cline-fitting analysis shows the steepness of the cline (Fig. 3c) and pinpoints the center of the hybrid zone to approximately 30–34 km south of Holbrook. The hybrid index analysis provides a visualization of the cline that also supports a sharp transition in population ancestry (Fig. 3a) and provides information about the relatively high abundance of late-stage backcross samples in comparison to F1 individuals (Fig. 3b). Finally, the introgression probabilities obtained for each hybrid population using the coalescent model illustrate the clinal nature of the hybrid zone by showing a clear pattern of increased introgression with geographic proximity with the largest change occurring between Canoncito and Sevenmile Draw (Table 4). The large shift in the genetic profiles of these populations is seen repeatedly by the shift in introgression probabilities measured with the MSC-I model (Table 4), the change in ancestry proportions (Table 1), and the cline-fitting and hybrid index analyses (Fig. 3). While these inferences are complimentary, the coalescent models add new information about the timing of introgression, the intensity of introgression from parental populations into hybrid populations, and the long-term effective migration rate between the parental populations.

### Coalescent analysis of the hybrid zone

Coalescent analyses of the *Sceloporus tristichus* hybrid zone using the MSC-I and MSC-M models provide new information about the evolutionary history of the study system, including estimates of the relative proportions of introgression into populations with mixed ancestry, the timing of their introgression, and the migration rates between the parental populations. The introgression probabilities (*φ*) show that the proportion of a hybrid population genome that is derived from a parent population increases as their geographic distance decreases (Table 4). This spatial pattern of introgression is consistent with the clinal nature of the hybrid zone, which shows a sharp transition in population ancestry over < 10 kilometers (Table 1).

Coalescent analyses of divergence times typically only provide estimates for speciation and/or population splitting event, while the MSC-I model adds a parameter for the timing of introgression upon secondary contact (Fig. 1). The estimated timing of secondary contact and introgression in the *S. tristichus* hybrid zone is approximately 10 kya, which is an order of magnitude younger than the divergence between the parental populations (Table 4). While it is possible that anthropogenic changes are currently influencing hybrid zone dynamics, the genealogical signal for introgression detected using the MSC-I model suggests a much older origination at the end of the Pleistocene (Table 4).

One limitation of the analysis of the MSC-I model is the assumption that introgression is constant among the sampled loci (Flouri *et al*. 2020), but differential selection is expected to cause introgression to vary across the genome. Genome-wide variability in introgression rates has been documented using the MSC-I model in *Anopheles* mosquitoes (Flouri *et al*. 2023) and *Heliconius* butterflies (Thawornwattana *et al*. 2023), which both show considerable variation in introgression rates across chromosome arms. The ddRADseq loci analyzed here are too sparsely sampled to describe the patterns of differential introgression that are likely to occur across the *S. tristichus* genome. One region of the *S. tristichus* genome that is of interest for a survey of variable introgression is the large pericentric inversion polymorphism on chromosome seven (Bedoya & Leaché 2021). Chromosomal inversions can gain a selective advantage if they capture two or more alleles that are locally adapted to the environment (Faria *et al*. 2019). Recombination suppression underlies this mechanism, because when locally adapted alleles are located within an inversion, they are no longer able to recombine with maladapted alleles located outside of the inversion (Kirkpatrick 2010).

It is important to consider how the assumptions of the MSC-I model could influence our interpretation of the hybrid zone. The simplified model that we used for the MSC-I analyses only includes two parental and one hybrid population (Fig. 1a), which makes it simple to implement and easy to interpret. More hybrid populations could be added to the analysis, but this would greatly increase the number of model parameters and make it difficult to obtain reliable estimates. It is possible to estimate the rates and directions of migration events between multiple hybridizing populations using MIGRATE-N (Beerli & Felsenstein 2001), but this model cannot account for the phylogeny or the history of population divergence events. The phylogenetic framework used in the MSC-I model enables the estimation of divergence times and introgression times. Finally, the introgression probabilities estimated by the MSC-I model assume a single pulse of introgression at the time of secondary contact. An alternative model that assumes constant migration throughout the duration of the population history is implemented in the MSC-M model, but this model does not estimate the timing of secondary contact (Fluori *et al*. 2023). A previous study of *Anopheles* mosquitoes suggested that the introgression events detected by these models may be largely historical, and that the MSC-I model could be closer to reality compared to the MSC-M model in cases where introgression varies through time (Fluori *et al*. 2023).

### Hybrid zone dynamics

We describe the clinal nature of a moving hybrid zone in *Sceloporus tristichus* using three replicate transect samples collected over two decades. In many ways, the characteristics of this hybrid zone fit the common tension zone model (Key 1968), which assumes that hybrid zones are maintained by a balance between selection against hybrids and dispersal into the hybrid zone and that parentals have higher fitness than hybrids (Barton & Hewitt 1985). The *Sceloporus* hybrid zone is situated at a habitat ecotone, which, while not required by the tension zone model, is true for some presumed tension zones (Endler 1977). When a hybrid zone is restricted to an ecotone, any environmental or climatic changes that result in movement of the ecotone could cause a shift in the location of the hybrid zone (Taylor *et al*. 2015). Hybrid zones have been documented to move in numerous plant and animal species at both contemporary (Buggs 2007) and historical times (Wielstra 2019). A unique benefit of the *S. tristichus* study system is the ability to leverage temporal sampling to measure how the center and width of the hybrid zone changes through time (Table 3). Replicate sampling of the hybrid zone every decade suggests that the center of the cline is shifting to the north despite the relative short duration between sampling intervals (10 years = ∼10 generations), but these small shifts in cline center are not significant (Table 3).

A null model for any hybrid zone is neutral diffusion, in which there are no barriers to gene flow. As populations freely exchange genes, the cline widens over time as a function of dispersal rate. Here, we can use the estimates of the hybrid zone age to test the plausibility of this null model for the *S. tristichus* hybrid zone. Under neutral diffusion, cline width is given by:

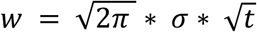

where *w* is the cline width, *σ* is the standard deviation of the distance between parents and their offspring, and *t* is the time since secondary contact (Endler 1977). We can take time since secondary contact as *t_hybridization_*, as reported in Table 4 as ∼11,000 years or ∼11,000 lizard generations and *σ* as 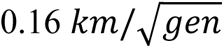 (inferred from a congeneric species *S. grammicus*; Sites *et al*. 1995). Thus, under a model of neutral diffusion, we would predict a cline width of 40 km – more than four times greater than what we see in this system (cline width ∼9 km; Table 3).

Because neutral diffusion is improbable, the hybrid zone between *S. tristichus* lineages is more likely maintained by environmentally mediated selection and/or selection against hybrids (Endler 1977; Barton & Gale 1993). The *S. tristichus* lineages occupy distinct habitats and have corresponding eco-morphologies – the northern and southern lineages are found in woodlands and grasslands respectively, and the southern lineage is both smaller and darker than the northern lineage (Leaché & Cole 2007). The hybrid zone between these lineages maps to the ecotone between them. Together, this evidence suggests that the hybrid zone is likely partially maintained by environmentally mediated selection.

We can use the predictions from the tension zone model to better understand the plausibility that the hybrid zone is maintained by selection against hybrids. Given cline width and a measure of dispersal, we can estimate the strength against hybrids using the equation:

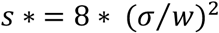

Again using the *σ* estimate from *S. grammicus*, and estimating the average nuclear cline width at 9 km (Table 3), we infer selection against hybrids to quite weak - just 0.2%. However, chromosome 7 contains a large pericentric inversion between the two lineages. While the role this inversion plays in local adaptation or reproductive barriers is yet unclear, cline width at the inversion is much narrower than at the rest of the nuclear genome (1.5 km vs. 9 km; Table 3; Leaché & Cole 2007). Selection on these inversions is predicted to be stronger (∼9%). Thus, if selection against hybrids is maintaining this hybrid zone, this selection is likely relatively weak and heterogeneous across the genome.

Whatever forces are maintaining this hybrid zone, they are strong enough to preserve the genetic integrity of the hybridizing lineages. Inside the hybrid zone, our results suggest rampant hybridization and introgression (Table 4). Outside of the hybrid zone, the outermost sampled populations are separated by 60 kilometers and show very limited levels of gene flow (*N_e_m* ∼ 0.03 and 0.3; Table 5). This hybrid zone thus acts both as a geographic border between these two lineages and a relatively impermeable barrier to gene flow (McEntee *et al*. 2020).

What then do these results tell us about the species status of the two *S. tristichus* lineages? The two lineages of *S. tristichus* that meet in the hybrid zone are estimated to have diverged ∼250 kya (Table 5), which is much younger than the multimillion-year divergences seen among recognized species in the group (Leache *et al*. 2016). Further, although the lineages differ at multiple inversions and their mitochondrial genomes are quite divergent (∼10% different), they only have modest levels of genetic differentiation (mean F_ST_ = 0.09; weighted F_ST_ = 0.23). Thus, despite their differing ecomorphology and some evidence of reproductive barriers between them, the two lineages appear to be relatively early on their journey to being robust, independent species.

Mitochondrial DNA introgression is not uncommon in hybrid zones, but a unique feature of the *S. tristichus* hybrid zone is the co-occurrence of four distinct haplotypes. First, the northern and southern haplotypes of *S. tristichus* form a clinal pattern with the major transition in their frequencies occurring near the center of the hybrid zone between Canoncito and Sevenmile Draw (Table 2). Previous studies of the hybrid zone have demonstrated that the mtDNA cline lags behind the nuclear cline by approximately 3 km (Leaché *et al*. 2017). However, haplotypes from two other clades enter the hybrid zone, including western *S. tristichus* haplotypes that are found in low frequency and *S. cowlesi* haplotypes that enter from the east and are in relatively high frequency throughout the hybrid zone (Table 2). Neither of these two haplotypes (western *S. tristichus* or *S. cowlesi*) have been detected in the pure parental populations.

Discordance between mtDNA and nuclear loci, either at the level of discordant spatial clines or conflicting phylogenies, is important to describe since these data are often used to delimit species. For example, the species that are interacting in this hybrid zone, *S. cowlesi* and *S. tristichus,* were delimited using mtDNA (Leaché & Reeder 2002); however, mtDNA introgression has resulted in spatial patterns that are distinctly different from population boundaries estimated from nuclear data. Detailed analyses of nuclear data are needed to clarify the species boundaries between *S. tristichus* and *S. cowlesi* and to determine the full extent of mtDNA introgression.

## Acknowledgements

We thank A. Carvalho, S. Wikramanayake, L. N’Diaye for help with fieldwork. We thank x anonymous reviews and the AE for their helpful comments and suggestions. We thank for the Arizona Game and Fish Department for issuing scientific collecting permits (LIC #SP843452). This project was supported by grants to A.D.L. (NSF-SBS-2023723) and S.S. (NSF-SBS-2023979).

## Notes

### Competing Interest Statement

The authors have declared no competing interest.

